# Bayesian Inference Framework to Identify Skin Material Properties *in vivo* from Active Membranes

**DOI:** 10.1101/2025.11.11.687900

**Authors:** Mark Wilkinson, Khushal Goparaju, Laura Nunez-Alvarez, Craig J. Goergen, Andres F. Arrieta, Adrian Buganza Tepole

## Abstract

Accurate *in vivo* characterization of skin mechanical properties is essential for diagnostics and treatment planning across dermatological and surgical applications. Existing noninvasive techniques are limited in capturing the nonlinear and anisotropic behavior of skin. In this work, we propose a Bayesian inference framework that leverages active membranes to induce desired deformations and infer patient-specific skin properties from a measured strain field. A finite element model of skin-membrane interaction, parameterized using the Holzapfel-Gasser-Ogden model, is used to generate strain field data under various membrane actuation conditions. To overcome the computational cost of repeated simulations required for Bayesian sampling, we construct a data-driven surrogate using principal component analysis for dimensionality reduction and Gaussian process regression for rapid evaluation. Our approach enables probabilistic inference of key skin parameters, including shear modulus, fiber stiffness, dispersion, and orientation. An advatange of the proposed method is that inference of skin biomechanics does not require direct force measurements. Rather, the method relies on known properties of active membranes (which can be tested ahead of time). The method does require strain field measurements. Through synthetic studies, we demonstrate that our method accurately recovers most model parameters even under moderate levels of spatially correlated noise, and that multi-frame or multi-membrane observations significantly enhance identifiability. These results establish the potential of active membranes as a viable platform for noninvasive, *in vivo* skin biomechanics assessment.

## 1. Introduction

Accurate measurement of skin mechanical properties is crucial for understanding disease and optimizing treatment for a wide range of pathologies. For example, hypertrophic dermal scarring, which can occur after severe burns or trauma, is characterized by increased stiffness [1, 2]. Treatment options include applying pressure on the tissue to gradually remodel the microstructure and promote a change in tissue-level mechanical properties [3], laser treatment to locally damage the tissue and induce its remodeling [4], or surgical revision [5]. Establishing an understanding of skin biomechanics is crucial for reconstructive surgery planning beyond scar revision. For instance, excessive mechanical stress at sutures causes an acute rise in inflammatory signals that can lead to complications such as delay healing and scarring [6, 7]. Thus, anticipating stress distribution in skin from reconstructive surgery to avoid complications requires establishing methods and experiments to determine the mechanical properties of skin [8, 9]. Diagnosis of skin cancer or monitoring skin diseases such as psoriasis can be done through the measurement of skin biomechanics [10, 11, 12]. Aging also leads to changes in skin biomechanics that can make it more prone to common aging features such as wrinkles [13, 14, 15].

Detailed mechanical characterization of skin can be easily conducted with established *ex vivo* testing protocols such as uniaxial tension, biaxial tension, or bulge testing [16, 17, 18]. However, this type of test requires excision of skin tissue samples, which is not always possible. To address this need, there have been numerous attempts to measure skin biomechanics noninvasively *in vivo*. The most successful noninvasive methods for testing skin mechanically include shear wave elastography and suction tests [19, 20]. Indentation of skin has also been attempted but the nonlinear deformation and involvement of underlying tissues makes it difficult to extract skin properties from these tests [21]. Flynn *et al*. developed a method to test skin *in vivo* with a micro-manipulator that loads the skin in various modes while recording force [22]. Simple methods such as shear wave elastography and suction are reliable, yet, only allow for determination of skin stiffness in the linear, small deformation regime [1, 23]. The micro-manipulator approach [22], can be used to probe the nonlinear response of skin, but requires expensive experimental setup.

Skin is a highly nonlinear and anisotropic tissue, routinely subjected to large deformations [24, 25]. Thus, even though linear approximations to its mechanical behavior are useful in the medical applications mentioned above, these approximations are many times too limiting [9]. Current noninvasive tests such as suction and shear wave elastography do not provide a rich enough set of deformations to infer the fully nonlinear behavior of skin [26]. Even though complex experimental setups such as the micro-manipulator proposed by Flynn *et al*. are up to the task [22], they are not easily deployable at a large scale to the general public.

Active membranes that can be applied to skin have been developed in the last decade. Initial work by Wong and co-workers [7, 27], showed that pre-stressed membranes could be applied across incisional wounds as a form of stress-shielding, reducing scarring. The work has been extended to different kinds of wounds such as chronic wounds and diabetic ulcers [28, 29], whereby a programmed strain on the membrane is used to drive wound contraction and promote healing. Pre-stressed membranes have recently been explored as cosmetic tools to increase skin pre-strain *in vivo* [30, 31]. Therefore, there exists technologies to apply desired strain fields on skin with pre-stressed membranes. These membranes can produce strains on the order of up to ten percent [30].

Techniques such as suction tests and shear wave elastography often assume a linear mechanical response of skin, enabling straightforward solutions to the inverse problem of estimating the skin modulus from measured variables [1, 19]. For nonlinear models of skin, analytical solutions for skin stress as a function of deformation are only available for cases of simple deformation [32, 25]. For complex deformation, e.g. the rich set of deformations from micro-manipulators or indenters [22, 33], methods such as inverse finite elements are needed [34]. If full-field deformation data is available, for example through digital image correlation (DIC), popular inverse methods can be used, including virtual element method and finite element updating [35]. A key limitation of these deterministic methods is that they are typically posed as least-squares optimization problems, despite the inverse problem being fundamentally ill-posed due to the significant inter-individual variability, non-uniqueness, and measurement uncertainty in skin mechanical properties. [9, 36]. Bayesian approaches are the natural alternative to infer probabilistic material parameters from complex deformation data [26]. A limitation of Bayesian methods, on the other hand, is the need for hundreds of thousands of forward model evaluations. To circumvent this limitation, we and others have developed data-driven surrogate models to replace expensive finite element simulations of skin [37, 38]. In this work we construct lower-dimensional representations of full-field strain using principal component analysis (PCA) and pose the inverse problem as a Bayesian inference problem in this reduced space.

In this manuscript we numerically investigate the potential use of *active* membranes as a means of identifying the nonlinear mechanical properties of skin noninvasively. Our results suggest that such programmable loading systems, combined with full-field deformation data and probabilistic inference, offer a promising strategy for patient-specific skin characterization.

## 2. Materials and methods

### 2.1. Forward model

We first model the skin and surrounding tissue and its interaction with an active membrane as the basis of our data generation. Figure 1 shows a visualization of the tissue model and the actuating membrane deformation. Here, the skin is modeled as a single thin homogeneous layer with underlying muscle tissue below. A 50 × 50 mm^2^ rectangular section was created in the finite element package ABAQUS, with section thicknesses for the skin and muscle being 0.6 mm and 10 mm respectively. Both layers are assumed to have only a hyperelastic response in a quasi-static simulation. The skin section is modeled with the Holzapfel-Gasser-Ogden (HGO) strain energy density function [39] as it has been widely adopted to model a variety of collagenous tissue, including skin [32, 17]. A key feature of this model is the anisotropic component which accounts for collagen fiber orientation and can contribute significantly to the mechanical response of tissue under high strain. The strain energy density has an additive split and takes the form

**Figure 1:**
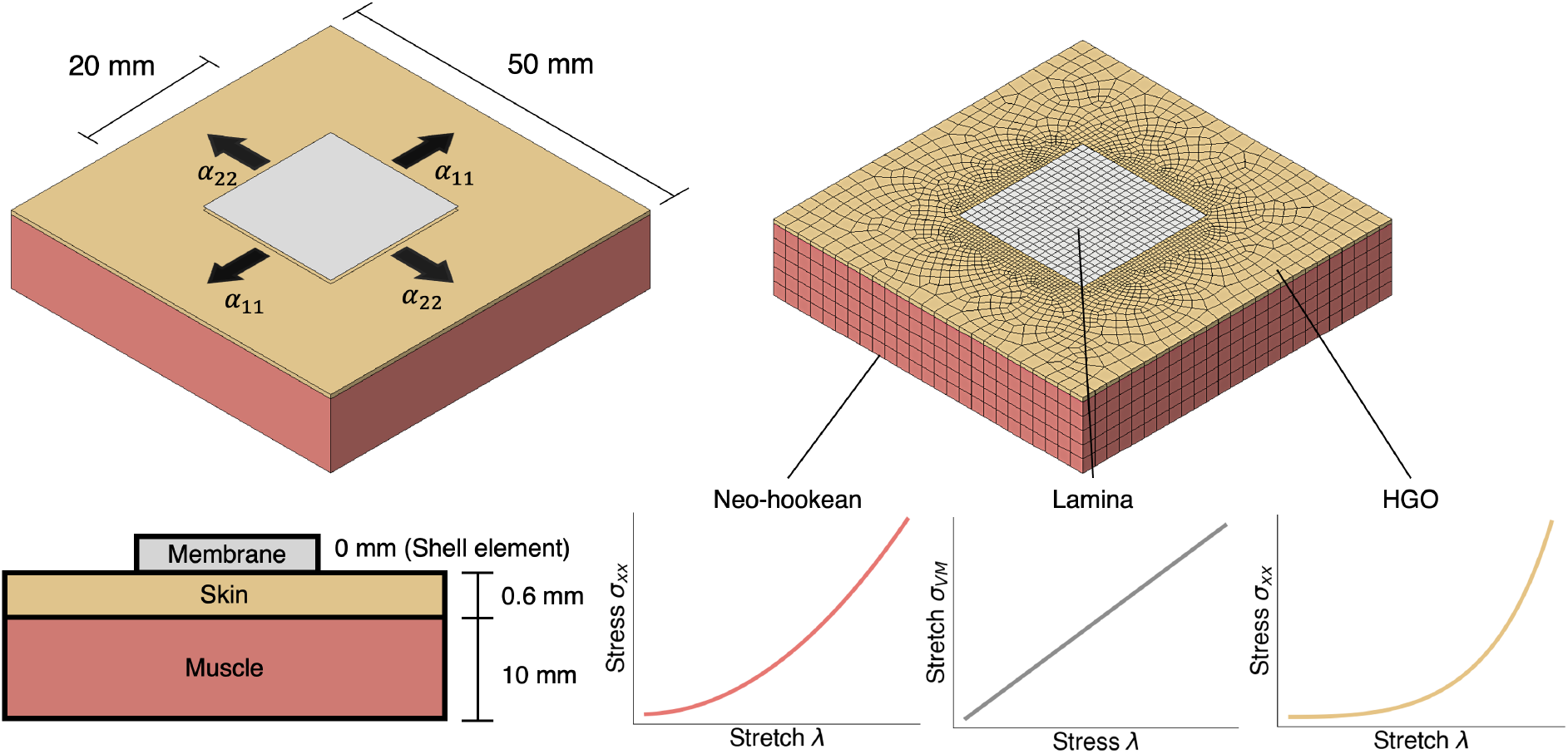
Finite element simulation of a representative tissue section. A 50 × 50mm^2^ tissue composed of skin and muscle layers was modeled with an attached linear elastic membrane on top actuated to deform in plane with eigen-strains *α*_11_, *α*_22_. Muscle was modeled as neo-Hookean, skin based on the HGO strain energy, and top membrane as linear elastic shell or *lamina* in Abaqus.

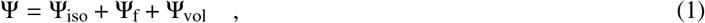

where Ψ_iso_ is the isotropic *ground matrix* component, Ψ_f_ is the contribution of the anisotropic collagen fibers, and Ψ_vol_ is a volumetric term. The functional form of the skin strain energy density function is

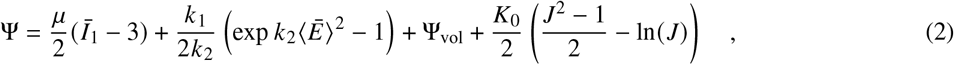

where the first term corresponds to Ψ_*iso*_ and it is the neo-Hookean ground matrix contribution specified by the shear modulus *μ*. *k*_1_, *k*_2_, and *κ* are anisotropic material parameters. Ē is defined as

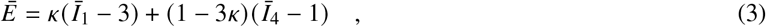

where Ī_1_ is the first invariant of the isochoric right Cauchy deformation tensor, 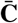, and 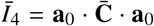 is the pseudo invariant with respect to the collagen fiber orientation unit vector **a**_0_. Additionally our volumetric term depends on the bulk modulus *K*_0_ and we take the ABAQUS default recommendation for this parameter with *K*_0_ / *μ* = 20. Note that we have consider a single fiber direction *N* = 1 in this model. Therefore the HGO model is fully parameterized by *μ, k*_1_, *k*_2_, *κ*, and the direction of fiber orientation **a**_0_.

For modeling of the underlying muscle tissue, we use a neo-Hookean model using the parameters for the quadriceps muscle from Ref. [40]. We did not consider any parameter variation for the muscle tissue. In contrast, we considered that the skin properties were uncertain, with variation of the HGO parameters based on Ref. [17]. The model was meshed with an average element size of 2 mm and a refined mesh element size of 0.5 on the membrane-skin interface. Our model displays 22,488 C3D8H elements, with 3,748 elements composing the skin. A fixed boundary condition was applied to the bottom surface, with the sides of the skin and muscle having planar roller conditions. The skin starts in a stress-free state for simplicity but, as detailed below in the *sampling* subsection, the parameters were chosen to reflect the anticipated *in vivo* stiffness [17].

To simulate actuation, a membrane patch was created and coupled to the skin. The membrane is a 20 × 20 mm^2^ shell and is rigidly attached to the surface nodes of the skin. This membrane can expand and contract in plane given a prescribed eigen-strain (*α*_11_, *α*_22_) in both the *x* and *y* directions. The eigen-strain in this model is imposed as a thermal expansion in ABAQUS, but it represents the possible actuation due to magnetic loading [41, 42], or pre-stressed membranes [30, 31, 43]. To reach the final strain state, 10 time increments were imposed. Note that the simulations were considered quasi-static. The material model selected for the membrane was linearly elastic, though we do not analyze the stress response of the membrane itself. Membrane properties are based on the assumption of polydimethylsiloxane (PDMS) with embedded nickel flakes [42]. Considering a rule of mixtures and assuming a 5% volume ratio of nickel, the material constants for the shell element were selected as *E*_1_ = 10GPa, *E*_2_ = 5GPa, *v* = 0.481, *G*_12_ = 5GPa, *G*_13_ = *G*_23_ = 2GPa.

Though the goal of an actuated membrane is to being able to show auxetic properties via magnetic particles embedded in particular orientations [41, 42], pre-stressed polymer films commercially available can only produce contraction [30, 31, 43]. Thus, both types of deformation were considered, as well as intermediate cases. In this work we prescribe strains in the space of (*α*_11_, *α*_22_) ∈ [*−*0.05, 0.05]^2^, which includes fully auxetic, contractionary, and uniaxial designs.

### 2.2. Sampling, model order reduction, and Gaussian process surrogate

The goal of the present manuscript is to being able to solve the inverse problem of identifying skin material parameters from the observed deformation when the membrane attached to the skin is *actuated*. Posing this problem as a Bayesian inference problem, which is developed in section 2.3, requires thousands of forward solver evaluations to sample from the posterior distribution. It is infeasible to evaluate the finite element model that many times due to computational cost. Hence, we first replace the forward solver by a surrogate model. To build the surrogate we sample the material parameter space (the samples of material paramters are denoted **X**, evaluate the finite element model at these inputs, and obtain the strain data (denoted by **Y**). Gaussian process (GP) regression is then performed over this dataset. The GP evaluation takes on the order of milliseconds and enables sampling methods for the Bayesian inference problem [9, 37].

#### 2.2.1. Sampling methods

We consider variation of *in vivo* skin parameters based on Ref. [17]. The maximum and minimum values of the *in vivo* conditions are used to define bounds of uniform distribution for *μ, k*_1_, *k*_2_ and *κ* sampled via Latin Hypercube Sampling (LHS) [44]. Additionally, since there is a single family of fibers defined in the model, the orientation of the fibers in plane can be fully described by an orientation parameter *θ*. The angle *θ* is measured from the direction of stretch *α*_11_ in degrees. Due to the biaxial symmetry we consider *θ*∈ [0^°^, 90^°^]. Thus, our material model is fully defined by the set of parameters (*μ, k*_1_, *k*_2_, *κ, θ*) ∈ **x**. Table 1 shows the range of allowed samples for each parameter. LHS was performed over these values to generate 120 unique parameter inputs for our forward finite element model.

**Table 1:** Latin Hypercube Sampling (LHS) parameter ranges based on Ref. [17].

| | $\mu$ (MPa) | $k_1$ (MPa) | $k_2$ | $\kappa$ | $\theta$ ($^\circ$) |
| --- | --- | --- | --- | --- | --- |
| Maximum | $2.06 \times 10^{-3}$ | $2.25 \times 10^{-2}$ | 15.32 | 0.30 | 90 |
| Minimum | $5.43 \times 10^{-4}$ | $1.17 \times 10^{-2}$ | 8.31 | 0.27 | 0 |

The eigen-strain of the active membrane was sampled from [*−*5%, 5%] in 7 linearly spaced points. Instead of simulating all 49 possible combinations of (*α*_11_, *α*_22_), we first fixed *α*_22_ to either 5% or −5% and varied *α*_11_ over the range [*−*5%, 5%] with 7 linearly spaced points. Permuting all combinations of material parameters and actuations, 1,680 simulations were run and the logarithmic strain tensor, denoted **E** throughout the paper, of the skin elements were recorded for each time step. This resulted in our dataset **Y** ∈ ℝ^16,800×3,748^.

#### 2.2.2. Dimensionality reduction

The strain fields from the simulations are defined over 3,748 elements in the skin. However, we anticipate that the inherent the dimensionality is much lower [37, 8]. Therefore, we perform Principal Component Analysis (PCA) to find a reduced basis for regression. Our strain data, represented through **Y**, is of the form

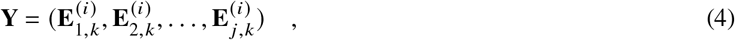

where 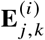 represents the strain tensor of a given element *j* at a integration point *k* in simulation *i*. We first average the strain across integration points such that we keep only one value of strain per element. We then split our data. We remove 20 randomly selected material parameter simulations from **Y** and denote this subset **Y**_*test*_. Then, we extract the *E*_*xx*_ and *E*_*yy*_ component of each element and split into two data matrices,

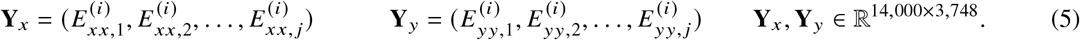

We center our data outputs by subtracting the element-wise mean and variance across each simulation to get

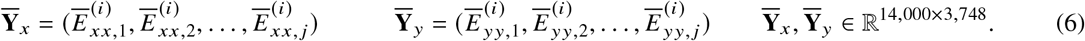

To begin the decomposition, we define **W**_*x*_, **W**_*y*_ such that

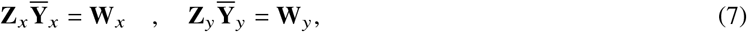

where **W**_*x*_, **W**_*y*_ ∈ ℝ^14,000×3,748^ are the principal component score matrices. To find each **W**, we do singular value decomposition on each matrix,

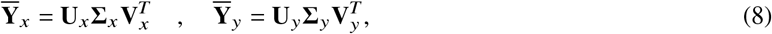

where **U, V**^*T*^ are orthogonal matrices and Σ is a diagonal matrix of non-negative singular values. For each system, we can rearrange the equations noting that since **U**^*−*1^ = **U**^*T*^, we can write

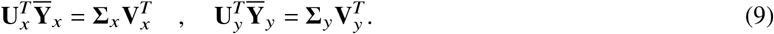

From this rearrangement, we can see that if we define **Z** = **U**^*T*^ and **W** = Σ**V**^*T*^ that we achieve the form of equation 7 for both components, where **Z** is the principal component vectors and **W** is the weight matrix. By performing singular value decomposition, the magnitude of the singular values show the most important features in our data. We can then represent our original data using a truncated matrix **W**^′^where **W**^′^∈ ℝ^14,000×*M*^, where *M* << 3, 748. *M* is chosen by finding the projection of the truncated PC basis back onto the original data such that 99% of the variance in the data is represented by *M* components. The test data **Y**_*x,test*_ can be also represented by a set of weights given by 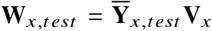 but using the PC basis of the training data. In other words, the dimensionality reduction is achieved based on training data alone, and **Y**_*test*_ is simply projected on the previously-identified reduced basis at the time of inference. Figure 2 shows a visual representation of PCA on our data with *M* = 8.

**Figure 2:**
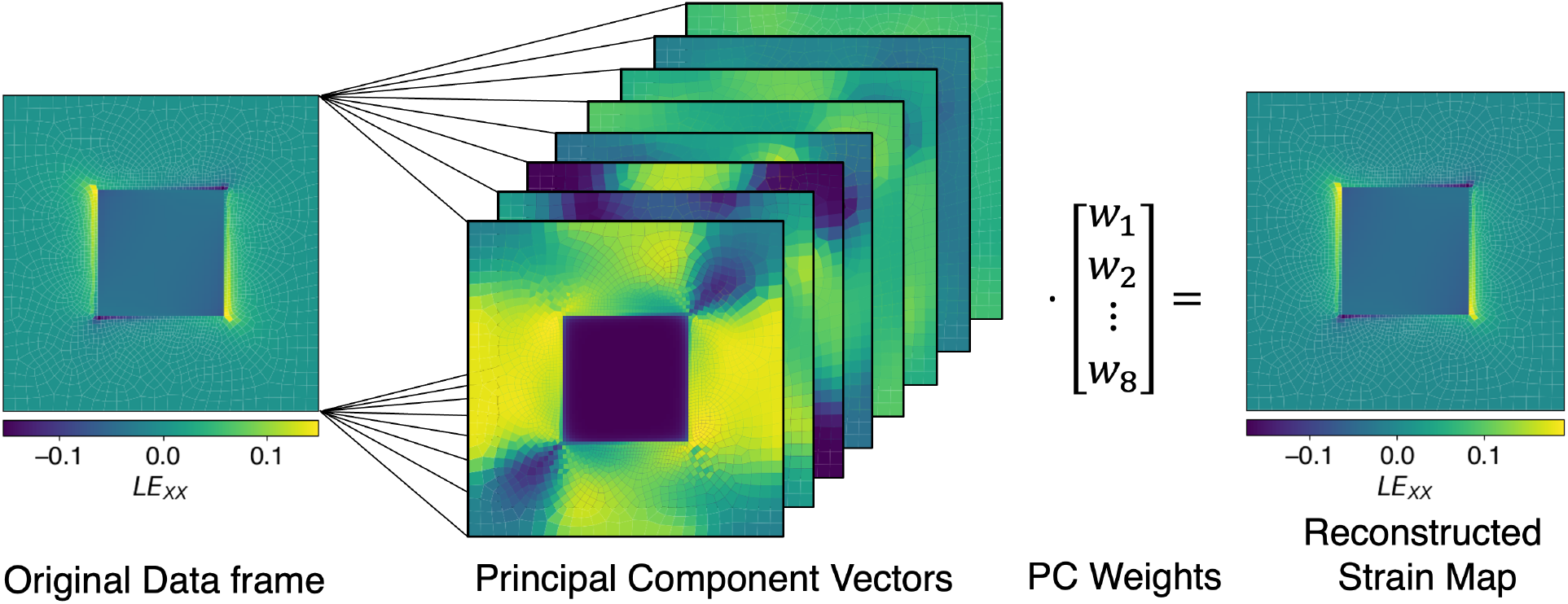
PCA decomposition of logarithmic strain *LE*_*XX*_ for the data. On the left is an original frame, which is then decomposed into a set of vectors and weights associated with each vector. This allows a reconstructed version of the data, shown on the right

#### 2.2.3. Gaussian process regression

Now having a reduced dimension for the targets, we define the input variables **x**_aug_ for regression as

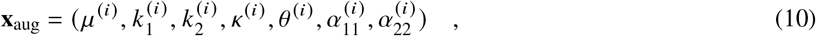

where 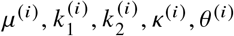 are the material parameters for a given simulation, and 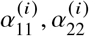 are the prescribed membrane strains. The targets of regression are the corresponding PC weights 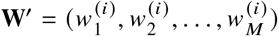. In other words, we want to learn the mapping between **x**_aug_ and **W**^′^.

We assume that the forward model outputs of the PC weights follow the form

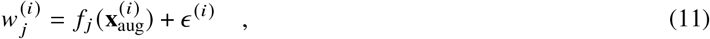

where 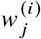 is the *j* th PC weight for a given simulation *i, f* _*j*_ (·) is a latent function for a given PC weight that we are trying to learn, 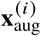 are the input variables for a given simulation, and 𝜖 ^(*i*)^ is an assumed noise to the observations. We take that *f* _*j*_ (·) is a Gaussian process and select the prior as

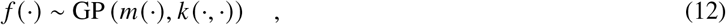

where *m*(·) is a specified mean function and *k*(·,·) is our covariance function. For this application we select a zero mean function and a radial basis for our covariance,

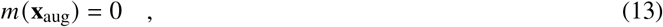

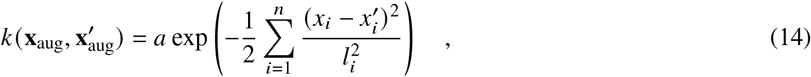

with 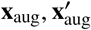 being two observations of our input space and 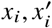 being a specific component of each of those input vectors. Here, the Gaussian process prior is controlled by the length scale hyper parameter *l*_*i*_ for each input, as well as the constant term *a*. This specifies the covariance matrix **K** where 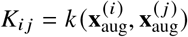.

To perform regression, a maximum likelihood estimate is sought for hyperparameter tuning to arrive at the poste-rior distribution 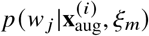, with *ξ*_*m*_ being the hyperparameters *l*_*i*_, *a*. The log-likelihood takes the form

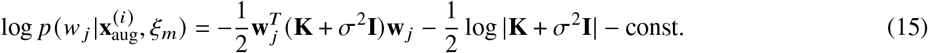

where *σ* is a specified noise added to the covariance matrix to stabilize the optimization. The const. term refers to constant terms which do not influence the optimization problem. After finding the hyperparameters through maximum likelihood estimation, we can find the posterior distribution of the PC weights, *w* _*j*_, which is another Gaussian process. Given some new input point 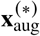, samples from the posterior can be taken from a normal over the function evaluations of new input points

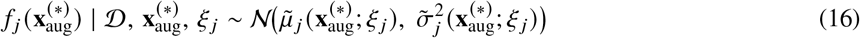

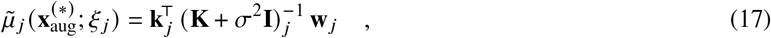

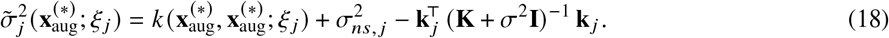

where 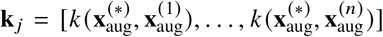 with the kernel being defined with the fitted hyperparameters *ξ* _*j*_ = (*a* _*j*_, *l*_1_, …, *l*_dim **x**_). Note that *w* _*j*_ is the *j* component of our PC weights, and the vectorization **w** _*j*_ in Eq. (17) comes from the multiple data points of that component.

This task was performed using Sci-kit-learn’s implementation of GP regression. For GP training, we further split our data. As described above, from the 120 skin parameters sampled, 20 were put aside for the test set and are thus used exclusively for inference of the inverse problem. The remaining data from the other 100 material parameters was used for PCA described above. Then, the PC wegiths are split into training and validation of the GP. Only 10% of the PC weight data was used in the GP covariance matrix as adding more data did not improve predictive capabilities and induced additional computational cost per iteration. GP’s are limited by the size of the covariance matrix **K** ∈ ℝ^*d*×*d*^ for *d* data points. Methods to use GP regression on large datasets, such as sparse GP’s [45], exist but were not used here for simplicity.

### 2.3. Bayesian inference for the inverse problem

To perform Bayesian inference, we follow the methodology proposed by Ref. [46], and show the approach in context the developed dataset and surrogate model. Let 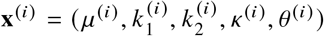be the material parameters, and let 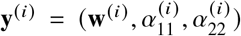 be the observations we have. We then assume that the parameters come from an existing probability distribution **x**^(*i*)^ *∼ p* (**x**^(*i*)^) known as the prior, and that our observations of the strain output are conditioned on the material parameters **y**^(*i*)^ |**x**^(*i*)^, **f**(**x**^(*i*)^) *∼ p* (**y**^(*i*)^ |**x**^(*i*)^, **f**(**x**^(*i*)^)) which is our likelihood given some model function **f**(·). This **f**(·) is our finite element model which approximates the observed data. The inverse problem is then formulated as sampling from the posterior distribution *p* (**x**^(*i*)^ |**y**^(*i*)^). Using Bayes rule, we have

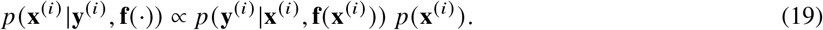

Evaluating this likelihood is difficult as it requires simulating the model for every sample of **x**^(*i*)^. Instead, a surrogate model can be used on the observations where we take a data set 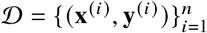with *n* training points. For some Bayesian regression task (such as the GP regression done in section 2.2.3), the predictive output **w**^*∗*^ given an input **x** is not deterministic. Therefore, the likelihood involves the sum rule of probability

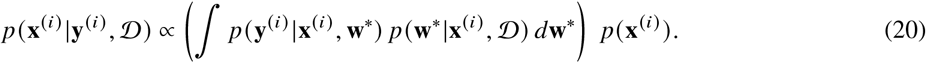

Intuitively, this corresponds to evaluating the GP surrogate at some input *p* (**w**^*∗*^ **x**^(*i*)^ 𝒟), but taking into account that the GP output **w**^*∗*^ is not actually deterministic. Instead, because **w**^*∗*^ is drawn from a Gaussian, this uncertainty needs to be marginalized or *integrated over*. We assume that all the priors are independent and have uniform distributions with bounds defined by the range of parameters sampled in Table 1. The prior is then specified by:

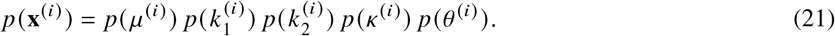

The likelihood is

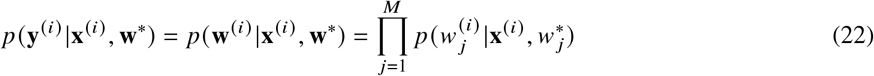

with *M* being the reduced basis dimensionality. Here we assume the likelihood is modeled as a normal distribution for each PC weight centered on the mean prediction *μ* _*j*_ (**x**^(*i*)^) of the surrogate model:

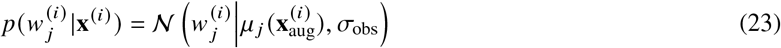

where *σ*_obs_ is a specified hyperprior and is given by a half-Cauchy distribution.

To sample the posterior distribution, we used Markov Chain Monte Carlo (MCMC) provided by the python package Pyro. We used the No-U-Turn-Sampler (NUTS) for our algorithm with an acceptance probability of 0.1 [47]. A single chain was run for 10,000 warm up steps and 10,000 samples. This was selected based on visual inspection of the chain producing uncorrelated samples and having reasonably smooth marginal distributions. For the likelihood, the forward model is evaluated from the sampled priors and the mean is taken from *p* (**w**^*∗*^ |**x**, 𝒟)as the forward prediction for a given input.

#### 2.3.1. Single observation

To evaluate the performance of the inverse problem, three different membranes are selected. These are (*α*_11_, *α*_22_) = (0.05, 0.05), (0.05, 0), and (0.05, *−*0.05). Each membrane is considered separately and only the last step of the simulation was considered, i.e. we assume that, for a given skin sample, we can only use a single membrane, and we can only observe the last frame of the deformation induced on the skin by that membrane. We evaluate the inverse model for the 20 skin parameters in the test set. We then compare the posterior distributions of the inferred parameters against the true parameters. Additionally, we find the 95% confidence interval and the mean of the posterior for a more quantitative comparison.

#### 2.3.2. Multi-observation

To further evaluate the effectiveness of this method, another probabilistic model was created. This model focused on incorporating multiple strain frames for a given skin sample out of the 20 in the test set. We anticipate that more observations should lead to increased predictive capabilities. Our prior remains the same, but we instead focus on providing multiple sets of PC weights **w**^(*i*,…,*n*)^ for *n* observations. These observations were selected through two different methods.

The first method involves providing data from multiple frames for the same simulation. This is equivalent to, given a membrane with actuation *α*_11_, *α*_22_, observing multiple frames as the membrane is gradually actuated over a given skin sample. For one simulation we have PC weights **w**^(1)^, **w**^(2)^, …, **w**^(10)^ for each of the 10 time steps. The inverse problem is run first with only the last time step and we add previous time steps one at a time to enrich information (i.e. (**w**^(10)^), which reduces to the single observation case, (**w**^(9)^, **w**^(10)^), (**w**^(8)^, **w**^(9)^, **w**^(10)^), etc.). MCMC was performed in the same way as before with this added information reflected in the likelihood.

The second method to increase the number of observations for the same material **x**^(*i*)^, involved having varying combinations of expansion parameters *α*_11_, *α*_22_. In other words, we assume that we could deform a given skin sample with multiple membranes, one after another. Only the PC weights of the last frame of the simulation are used for each membrane. The total number of membrane designs is 14.

#### 2.3.3. Impact of noise

To understand the impact of noise on the inverse problem, the original strain data **Y** had noise added to it via a Gaussian process which was not trained. This GP simply allowed us to generate spatially correlated noise to corrupt the strain fields. A zero mean and radial basis function were selected for the noise GP from equations 13 and 14 respectively. We investigated two different factors involved with noise generation. The first is the length scale of the basis function, where we selected a length scale of 0.1, 1, and 5 mm. Figure 3 describes the addition of noise at different length scales to the strain data. Additionally, the magnitude of the noise was adjusted. Samples of noise were initially from a standard normal distribution, and the magnitude was scaled by the percentage of the maximum strain given in the dataset for both the *x* and *y* strains. We investigated three different magnitudes of 1%, 5%, and 10% of the maximum strain. In combination with the length scale, there were 9 noise data sets created in total. These are labeled as a combination of “s”, “m”, and “l” for the length scale and “1”, “5”, and “10” for the noise magnitude. Therefore the data set “s5” refers to 5% noise magnitude with respect to the maximum strain over the entire simulation added at the smallest length scale. After generating, the same procedure for each dataset was followed as in the noiseless case by doing PCA, training a GP surrogate, and then performing Bayesian inference in both a single sample case and multiple samples.

**Figure 3:**
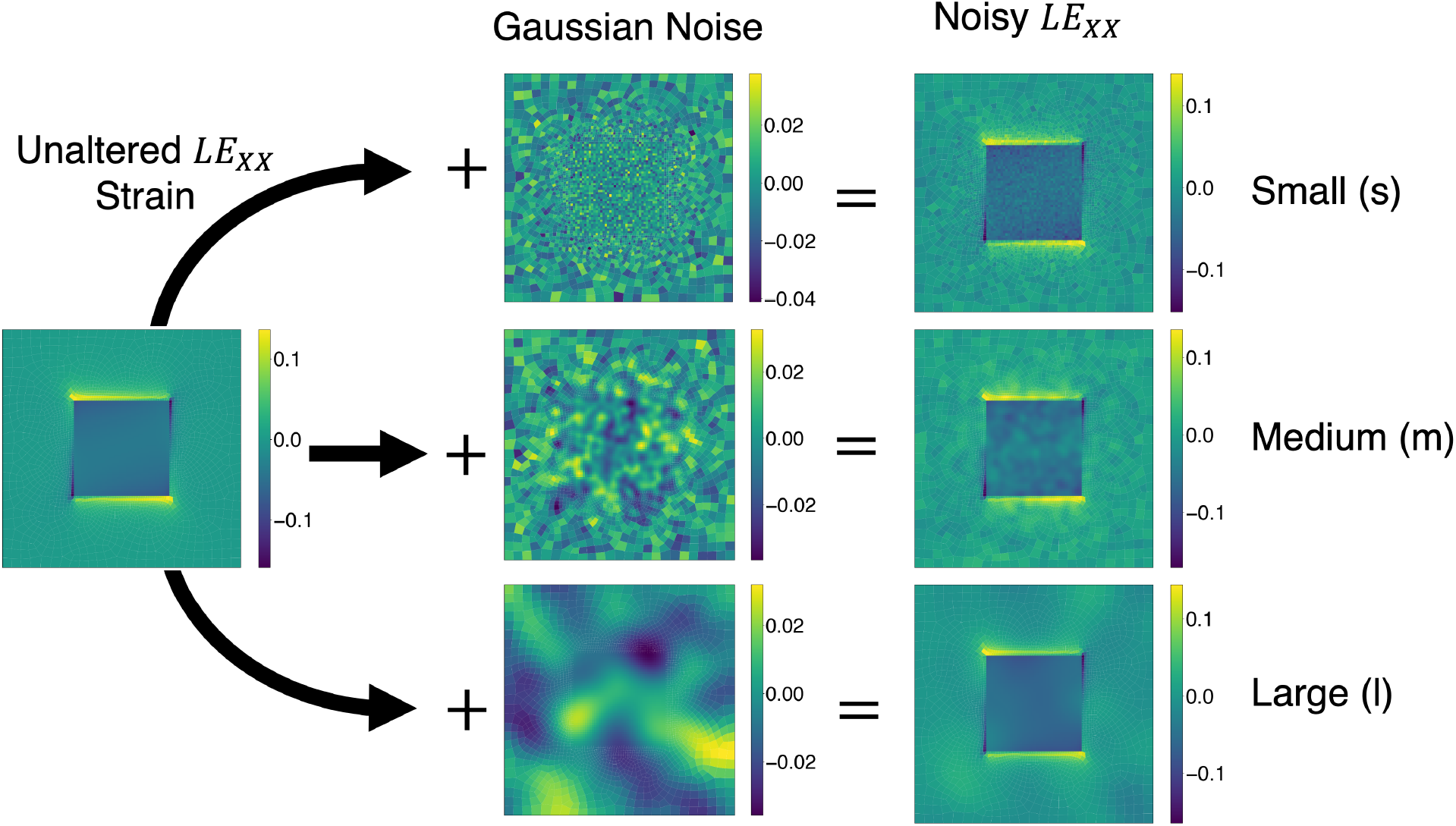
Corrupting strain field data with Guassian process noise. Guassian proccesses with zero-mean function and exponential kernel allow us to add spatially correlated noise to the strain field data with three different length scales shown. Noise magnitude was also varied in three levels:1%, 5%, and 10% of the maximum strain (not shown).

## 3. Results

### 3.1. Sampling and GP performance

As specified in the methods, 1,680 simulations were run in total, each containing 10 time frames. Data corresponding to 100 out of 120 material parameter samples 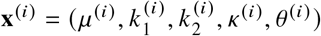and the corresponding strain fields **Y**∈ ℝ^14000×3748^, were first considered for PCA and then for GP training.

For the noiseless case, a threshold of 99% cumulative explained variance was decided as the cutoff point for the PC’s. This resulted in *M* = 8 components as our reduced basis. The decomposition is illustrated in Fig. 2 and the cumulative variance explained as a function of the number of PCs is illustrated in Fig. 4. This graph can represent the ability of the PC’s to reconstruct the data, with a cumulative variance corresponding to less error when projected back into the original data space. The choice of *M* = 8 is suitable output space for training the GP surrogate and involves only fitting 8 independent GP’s. However, PC reduction is sensitive to noise. Taking into account the spatially correlated noise in Fig. 3, PC performance drops drastically, as seen in Fig. 4. The magnitude of the noise increases reconstruction error significantly and is much more significant than the type of noise added. Note that the noise is added directly to strain values **Y** and is aimed at capturing the expected noise from experimental techniques such as digital image correlation (DIC). Over all the simulations and elements, the highest strain values were *E*_*x*_ = 0.2127 and *E*_*y*_ = 0.2777. Small noise denoted *s*1 corresponds to noise sampled from a GP with variance *σ* = 4.524 ×10^*−*6^ and thus the strain is perturbed by values in the confidence interval [*−*6.801× 10^*−*5^, 6.819 × 10^*−*5^]. With this level of noise the PC performance drops to around 90% at *M* = 8. A GP with variance *σ* = 1.131× 10^*−*4^ added to the strain data **Y** means we are perturbing the strain with values in the confidence interval [*−*3.400× 10^*−*4^, 3.409 × 10^*−*4^]. In this case, denoted *s*5, the PC performance drops to approximately 50% by *M* = 8 PC. Larger errors, considering perturbations to the strain by values in the confidence interval [*−*6.801 ×10^*−*4^, 6.819× 10^*−*4^] result in only < 30% of variance explained by the first 8 PC basis.

**Figure 4:**
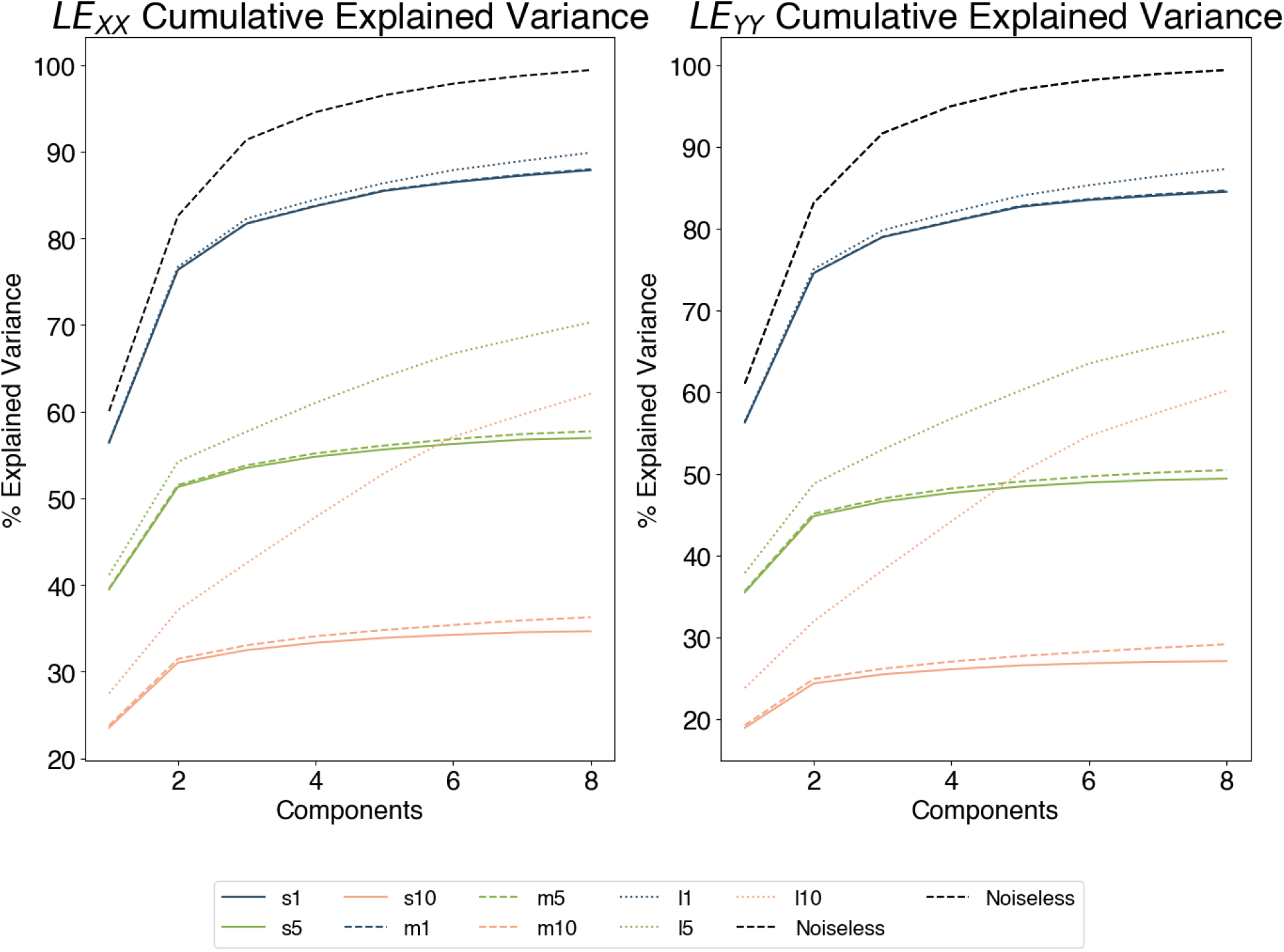
PCA cumulative variance explained for *x* and *y* strains. The noiseless data can be reconstructed with 99% cumulative variance explained using only the first *M* = 8 principal components. Spatially correlated noise in the strain fields leads to a drastic reduction in PC reconstruction. Noise magnitude is more detrimental than correlation length. 1% errors are well-tolerated, but 5% and 10% errors in strain lead to PC variance explained dropping to approximately 60% by *M* = 8. The effect of length-scale is less pronounced, with smaller length scale *s* leading to poorer performance than larger length scales *l*.

GP regression was performed on the reduced output separately for each noise level. In all cases *M* = 8 PC bases were used for comparison. As specified in the Methods, the GP was trained only on 10% of the PC data, with the remaining 90% used for GP validation. Figure 5 shows the accuracy plots of the PC weight by the GP surrogate for the noiseless case.

**Figure 5:**
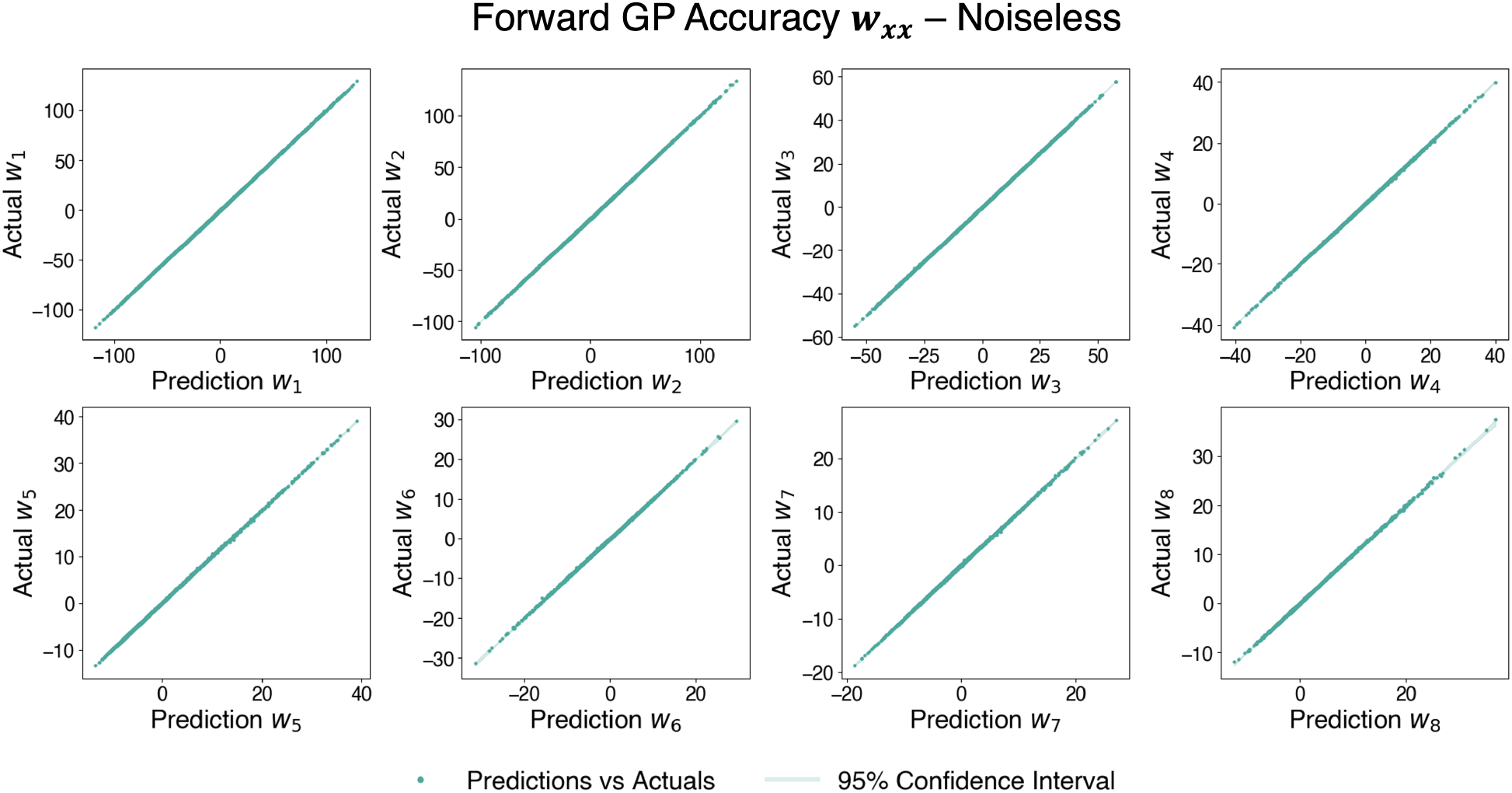
Accuracy plots for PC weights [*w*_1_, *w*_2_, …, *w*_8_] given the noiseless data. Each plot shows the GP output against the actual PC weights.

Tables 2 and 3 shows the *R*^2^ score for the noisy data sets. The length scale has much more dramatic negative impact on surrogate performance, as all the small length scale sets achieve a value greater than 0.9. Figure 6 highlights the performance for PC weight’s *w*_1_, *w*_5_, and *w*_8_ for different magnitudes of noise addition and spatial correlations. For all noise datasets considered here, the first component is the most accurately predicted while successive components lose predictive performance. The decrease in performance for predictive capabilities is related to how the components are related to the original data, since *w*_1_ is the best possible one-dimensional reconstruction of our data and contributes the greatest cumulative explained variance as can be seen in Figure 4. Successive terms add information to the reduced basis space with diminishing returns. Additionally, although larger spatial correlation data sets achieve better reconstruction when PCA was performed, forward surrogate performance is drastically reduced. We expect worse performance in Bayesian inference due to this loss in GP accuracy, though as will be shown later, there is still some information that can be extracted in the inverse problem even if the presence of large amounts of noise.

**Table 2:** *R*^2^ values for noisy *ε*_*xx*_ test set.

|  | 1% | 5% | 10% |
| --- | --- | --- | --- |
| Small | 0.99103 | 0.97237 | 0.94500 |
| Medium | 0.98351 | 0.84659 | 0.66895 |
| Large | 0.74527 | 0.32622 | 0.21678 |

**Table 3:** *R*^2^ values for noisy *ε*_*yy*_ test set.

|  | 1% | 5% | 10% |
| --- | --- | --- | --- |
| Small | 0.99004 | 0.96675 | 0.93292 |
| Medium | 0.97717 | 0.76137 | 0.57426 |
| Large | 0.59467 | 0.27261 | 0.19294 |

**Figure 6:**
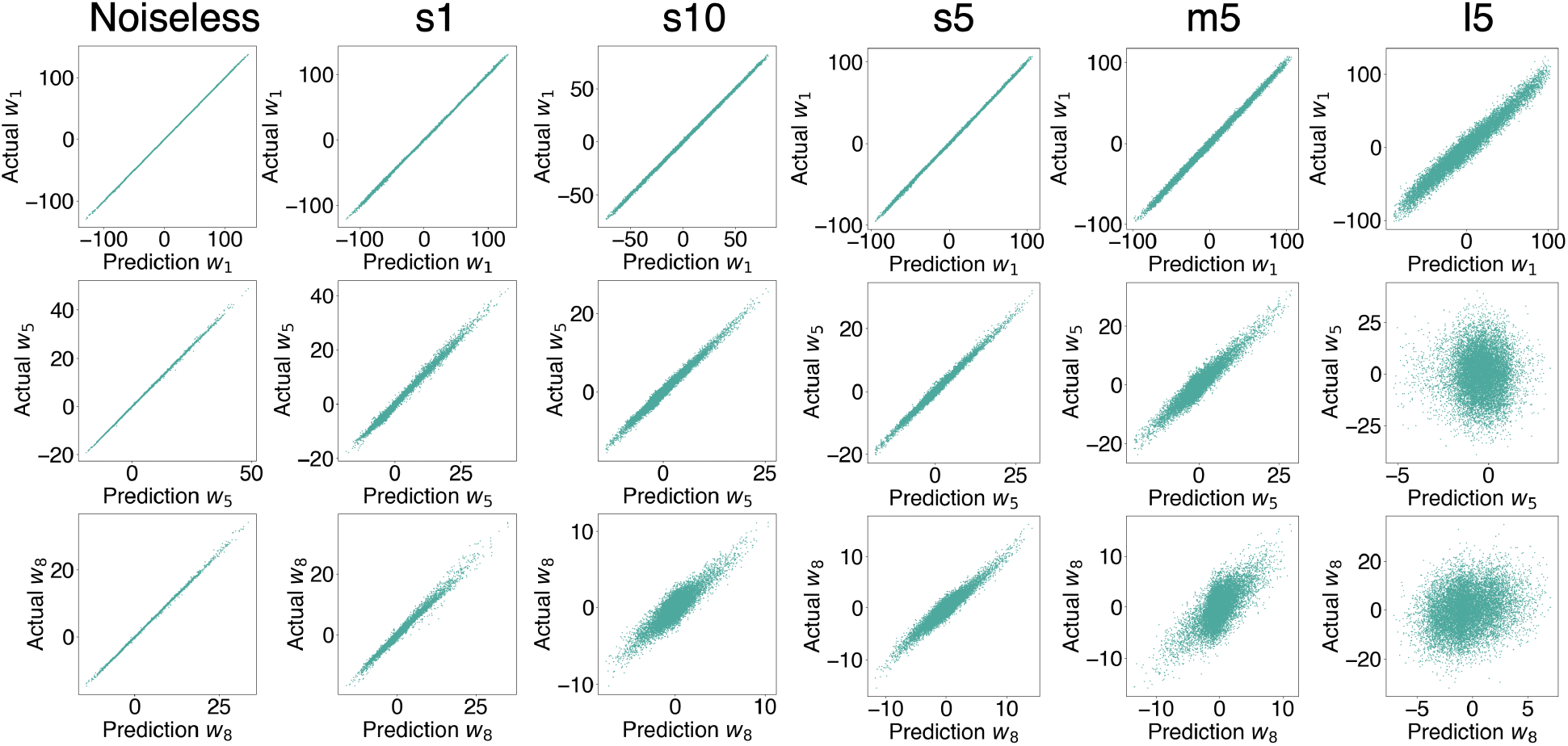
GP performance analysis considering both noise magnitude and length-scale variance. For the small length scale of noise, *s*, the addition of 1% and 10% noise leads to a slight drop in performance for the PC weights, with good performance for the most important weight *w*_1_ and increasing error for *w*_8_. For the same level of noise of 5%, varying the length between small *s*, medium *m*, or large *l*, had a more pronounced effect. The largest length scale led to significant error to the point that at weights *w*_5_ to *w*_8_ there is virtually no correlation between the predicted and observed PC weight.

### 3.2. Noiseless inverse problem

We recap that the inverse problem involves observing a strain field from the corresponding actuation of the membrane and the objective is to recover material properties of the skin. Even though we start with strain data **Y**, we then project it onto the PC basis to get the corresponding observations of the PC scores **w**. The observations for the inverse problem are thus the scores **w** together with actuation *α*_11_, *α*_22_ and the output of the inverse problem are parameters **x** = (*μ, k*_1_, *k*_2_, *κ, θ*). Below we show first, for noiseless training data, the effect of a single strain observation (or equivalent single PC vector observation **w**) on the inverse problem. Next we show that if we have more information, either because we have more frames for a given membrane or because we test the skin with different membranes, the posterior collapses more and our inference improves. Lastly we investigate the inverse model performance with the noisy cases.

#### 3.2.1. Single sample noiseless case

Figure 7 shows one out of the 20 material parameters that is in the test set (not used in PCA nor GP regression). The corner plot shows the posterior plots after MCMC sampling. Since our dataset it synthetic, we have the original values which generated a given simulation and thus the true values are signified by orange vertical lines in the diagonal of the corner plot (the marginal posteriors) or the orange dot in the pair plots in the off diagonal. Though this method is Bayesian in nature, we select the median as the dashed black line to compare against the true parameter as a point prediction. Off diagonal scatter plots show samples from our prior in orange and our posterior in green. The MCMC simulation is sampling in a 5 dimensional space and can’t be visualized directly, but using pair-plots we can see slices into this space and investigate how two parameters samples are related. The main outcome is that for the one case shown in Figure 7, we see the posteriors collapsing around the true parameters *μ, k*_1_, *κ* and *θ*. Posterior collapse is most evident in the case of the *θ* parameter which described the anisotropy orientation. The posterior is extremely narrow, centered on the true value of *θ*, and no uncertainty is apparent in this identification. In contrast, for *k*_2_ we see that our distribution does not collapse and instead remains closely uniform. This means that the PC observations do not contribute enough information to retrieve the true *k*_2_ value. The parameter *κ* is relatively well identified, but with a confidence interval [0.2885, 0.2988], which indicates that there is still some uncertainty in this parameter after calibration. The parameters related to moduli of background matrix and collagen fibers, *μ* and *k*_1_ respectively, are identified accurately with narrow posteriors *μ* :[0.00121, 0.00131] MPa, *k*_1_ : [0.0178, 0.0206] MPa.

**Figure 7:**
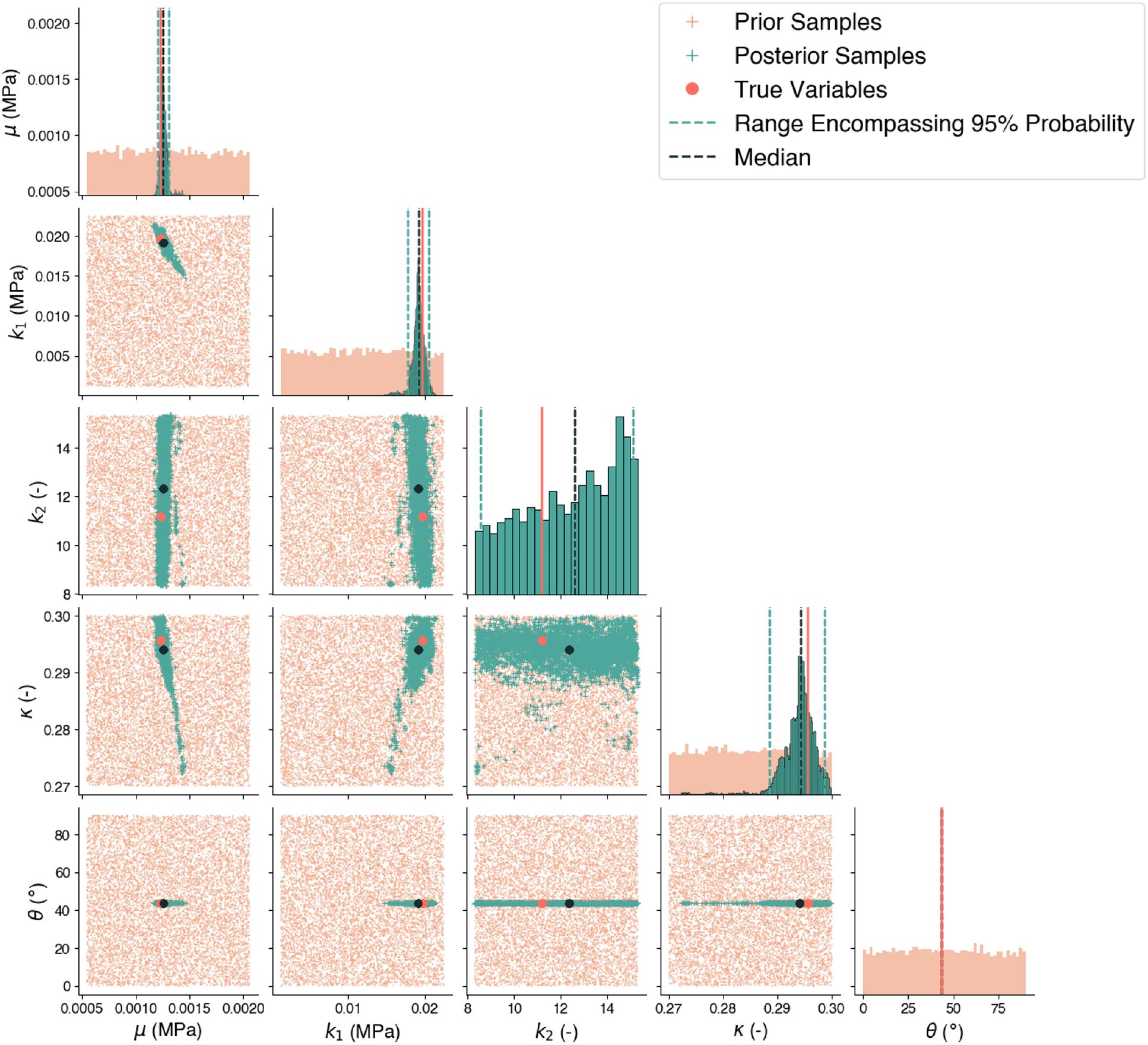
Inverse posteriors for the case of single noiseless observation. Prior distributions are plotted in orange and the posteriors in teal. Note that the variable *θ* collapses to extremely small window, while *k*_2_ appears to not collapse. The other material parameters, *μ, k*_1_, *κ*, are inferred accurately, with posteriors collapsing around the true value, but some residual uncertainty remains.

#### 3.2.2. Multi-sample noiseless case

Instead of providing a single observation **w**, we can instead provide multiple observations to enrich our inference. As specified in the Methods section, we select two types of multi-observation cases. First we use **w** coming from the last time frame of simulations with different expansions *α*_11_, *α*_22_. This would resemble using multiple different membrane designs experimentally as each would have a different prescribed deformation *α*_11_, *α*_22_. For the other case, we utilize multiple **w** coming from the same simulation but different time frames. This would be equivalent to having multiple strain maps extracted from the same membrane as it is gradually actuated. Figure 8 highlights these two data combination techniques and shows the result in the inference task. As expected more data leads to a narrowing posterior distribution. These results are directly explained by the fact that the likelihood function has more information provided to it, which better constraints the posterior samples. There are differences between the two multi-observation setups. The posterior collapse is similar in both cases with regards to the residual variance. The case with multiple membranes seems to better capture the true parameter values. In contrast, multiple frames for a single membrane design collapses the posterior but there is a discrepancy between the inferred and true parameters that is not alleviated by the addition of data. This would suggest that additional membranes is preferable to more frames from a given membrane. However, there could be other explanations, such as error in the GP near the boundary of the parameter range.

**Figure 8:**
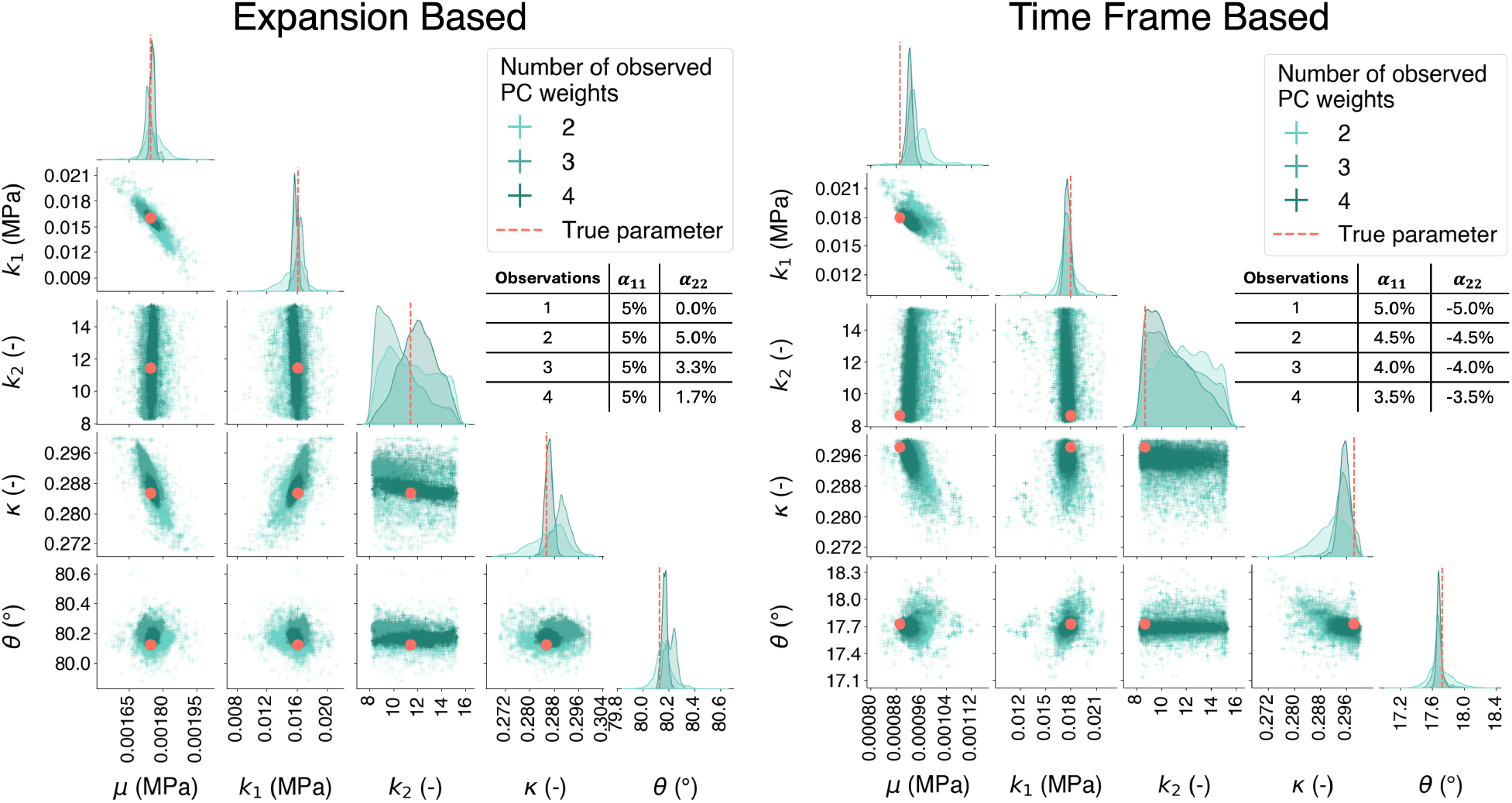
Comparison of the two multi-observation setups for the inference problem in the noiseless case. In one scenario, we assume that we can measure strain fields (or corresponding PC weights) from actuating the same skin (a single parameter set for skin) with different membrane designs (*α*_11_, *α*_22_ combinations). In the other case we assume that we can test a skin sample with only one membrane (a single *α*_11_, *α*_22_ final value), but we are able to record strain fields (or PC weights) gradually from no actuation (*α*_11_ = *α*_22_ = 0) to full actuation. In both cases, additional data leads to better posterior collapse, i.e., better inference. The performance in the multi-membrane case is better. In either case, all parameters except for *k*_2_ can be recovered.

One aspect that is consistent in both of the multi-observations setups is that *k*_2_ is not better inferred with respect to the single observation case. This implies that there is some information loss either in the forward model or in the PC representation of the data. Because PC in the noiseless case leads to 99 variance explained and the GP performance is excellent in the noiseless case, the loss of information about *k*_2_ is truly a feature of the forward problem. In other words, the idea of testing the skin with active membranes, even with various membranes, is not enough to recover *k*_2_. Despite this, all the other parameters are recoverable to a significant degree of certainty.

### 3.3. Inverse problem in the presence of noise

Figure 9 shows the inverse problem for the noiseless cases against the corrupted datasets for a mixed expansion regime of [*α*_11_, *α*_22_] = [5%, *−*5%] considering a single observation. The first row in Figure 9 shows all 20 test skin parameters in the case of noiseless data. These results align with the single example shown in Figure 7, which showed the correct identification of parameters for just a single skin material out of the 20 in the test set. The same trends from Figure 7 can be observed for all 20 test cases, which span the entire range of parameters. Some parameters are more discernible than others, with *θ* having the most accuracy in the inference, followed by *μ*, then *k*_1_, and *κ* still being identified but having a wider distribution. *k*_2_ is not discernible across the range of parameters, confirming that this issue is a feature of the forward problem. In other words, the strain fields do not contain enough information about *k*_2_, even in the noiseless case.

**Figure 9:**
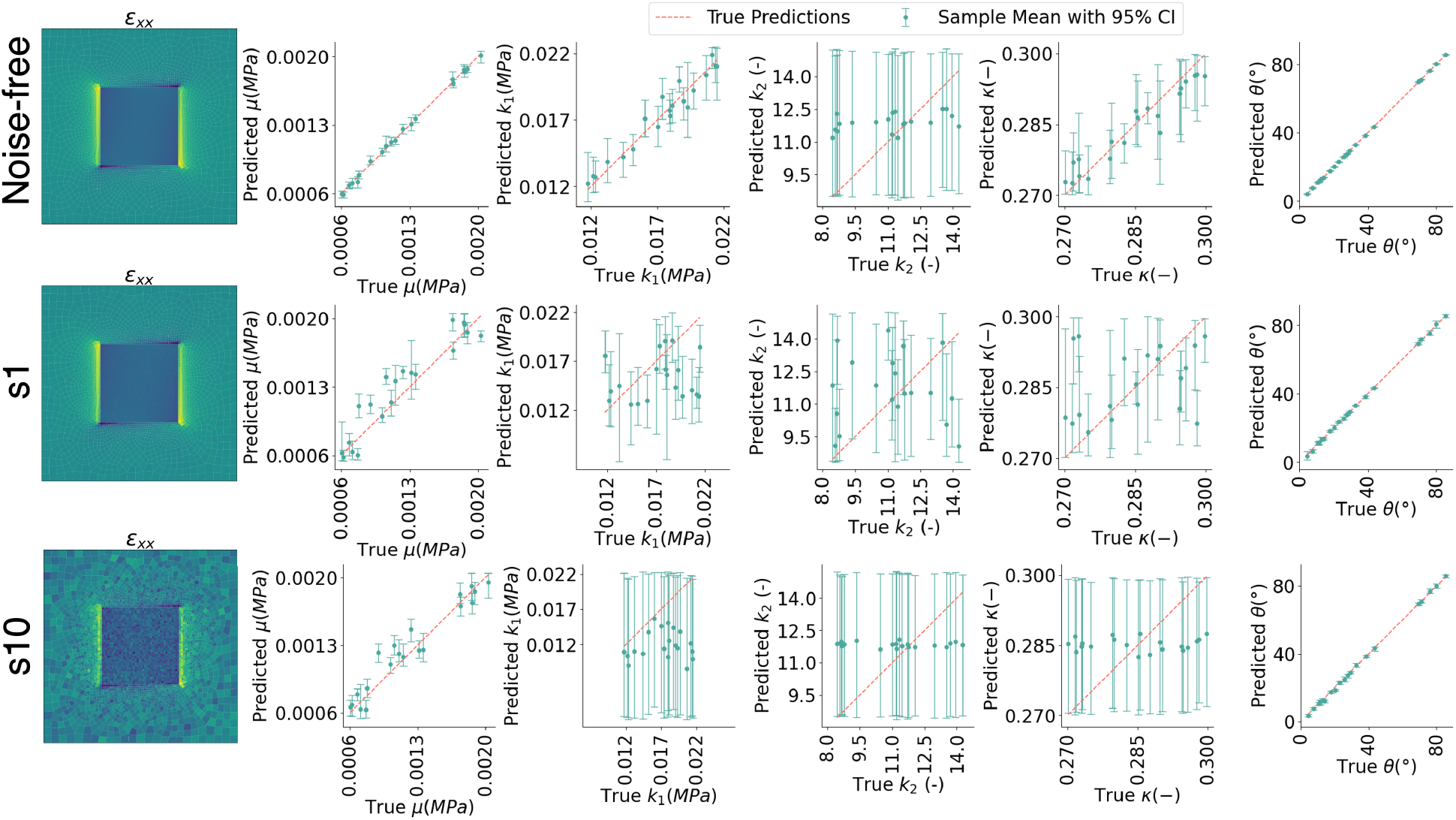
Performance of the inference problem across 20 test cases, in the noise-free and noisy cases. Example strain fields are shown. Accuracy plots show the comparison between the true parameter and the posterior of a MCMC run. In the noiseless case, all parameters but *k*_2_ are identified. As noise is injected, information about *κ* is lost first, followed by loss of identification of the *k*_1_ parameter. The shear modulus and anisotropy parameters, *μ, θ*, are identified well at all noise levels.

When injecting noise, there is an immediate affect on the inverse problem. The shear modulus and angle can still be identified with good confidence. *k*_2_ is still expected to perform poorly across all noise levels. *k*_1_ and *κ* are very sensitive to noise. Looking to the difference in performance in *k*_1_ from the s1 and s10 simulations, we see that there is still some information about *k*_1_ in the s1 case (which corresponds to errors on the order of 1% of the maximum strain). Yet, the performance in identifying *k*_1_ in the s1 case is considerably worse than in the noiseless case. s1 noise completely corrupts the inference of the dispersion parameter *κ*. At the highest level of noise, *s*10, *k*_1_, *k*_2_, *κ* are completely unrecoverable. However, *μ* and *θ* are still recoverable even at this level of noise.

To investigate the loss of predictive capability of *k*_1_, we analyzed the sensitivity of the PC components with respect to *k*_1_ and *μ* through the evaluation of the surrogate model. Figure 10 shows the output prediction of the first 4 PC components for the noiseless GP and the *s*1 GP. *μ* and *k*_1_ were sampled within the range specified in Table 1 while parameters *k*_2_, *κ*, and *θ* were kept constant at their mean value. The contour plots for the noiseless GP show that as *μ* is varied, the output of PC weights, **w**, significantly changes while *k*_1_ has a much smaller but still visible impact. When switching to the *s*1 case however, we see that the small influence of *k*_1_ almost completely disappears, i.e. the contours are nearly vertical which means that *k*_1_ has virtually no impact in the surrogate model prediction. Due to the high accuracy of the GP model established previously (Figure 5), we can say that this information loss is not due to the surrogate model but instead due to the underlying PCA representation, which is also supported by Figure 4. The information loss about *k*_1_ with the addition of noise is then reflected in a poorer identification of the *k*_1_ parameter during Bayesian inference.

**Figure 10:**
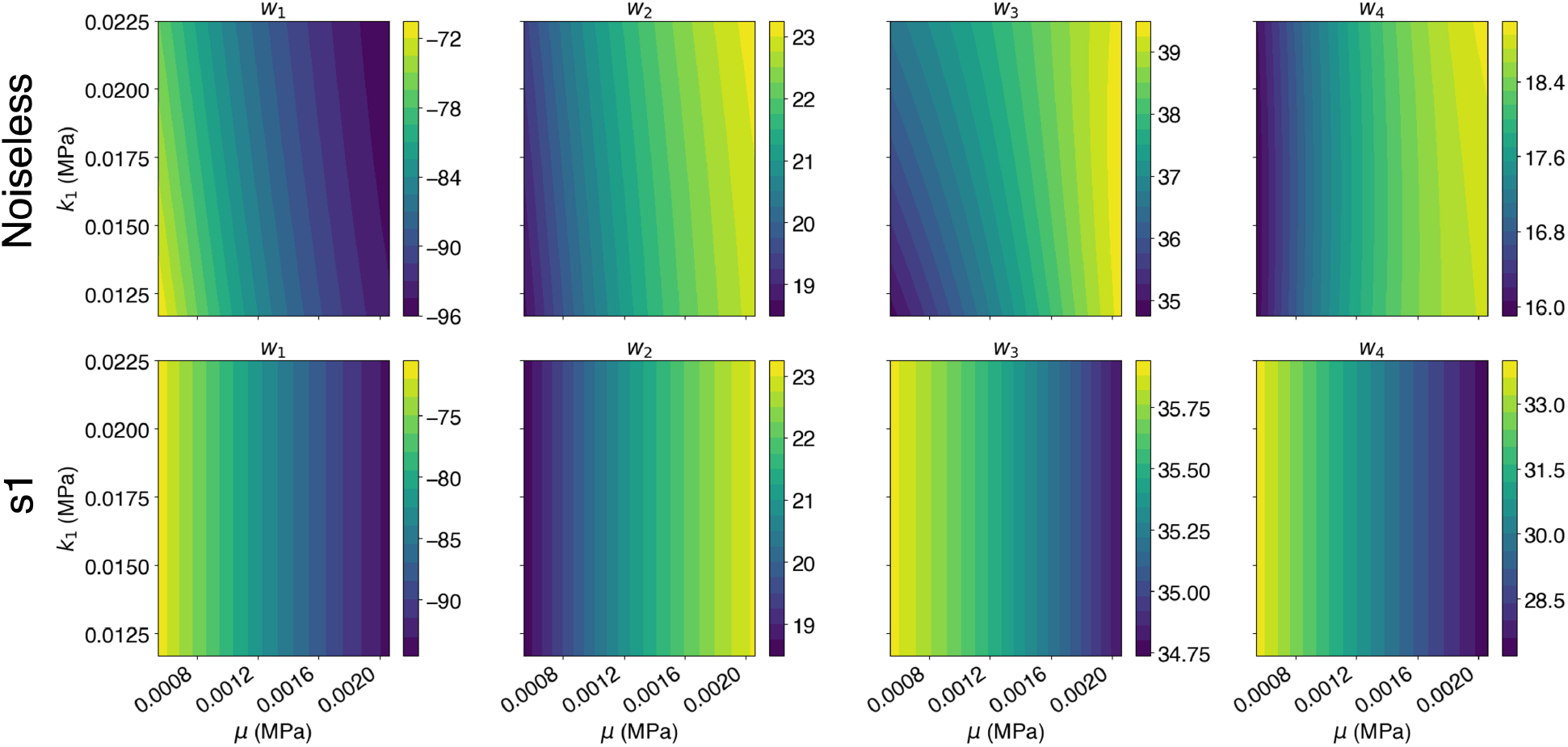
Sensitivity of principal component prediction through the surrogate model as a function of the *μ* and *k*_1_ parameters in the noiseless and s1-noise cases. Increasing noise renders the GP predictions insensitive to *k*_1_.

Figure 11 shows the effect of the length-scale of the spatially correlated noise described in Figure 3. As noise length scale increases, the performance of the inverse problem drops. The noise s5, which has a 0.1 mm length scale, experiences similar problems as discussed in Figure 9, with *k*_1_ and *κ* becoming more uncertain but *μ* and *θ* still recoverable with high accuracy. As we increase the length scale correlation of the noise, more information is lost even though the noise magnitude is kept at 5% of the maximum strain. When we corrupt the data with spatially correlated noise of large length scale in the correlation, *l*5, it becomes impossible to recover any of the parameters except *θ*. Note that this is also the only case shown where we have significant uncertainty in *θ*.

**Figure 11:**
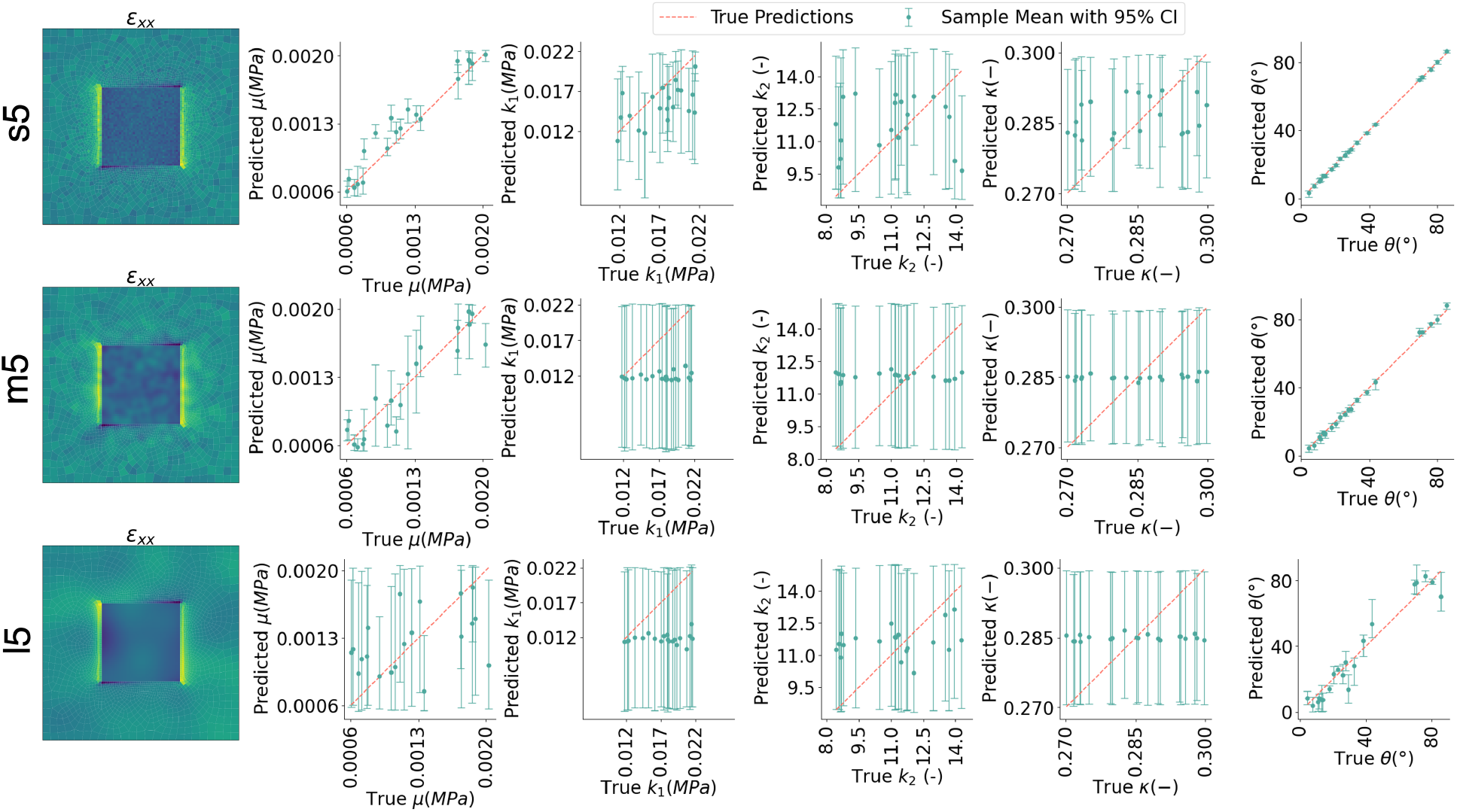
Effect of lenght scale for intermediate noise. These are given for [*α*_11_, *α*_22_] = [5%, *−*5%] single observation cases across all 20 of the test set skin samples. Increasing length scale in the noise leads to loss of information for all parameters.

## 4. Discussion

In this study, we proposed a Bayesian inverse problem to infer skin material properties from strain measurements obtained via actuated membranes attached to the skin. Our approach leverages a forward model of skin deformation under membrane actuation. Computational efficiency is achieved through dimensionality reduction using principal component analysis and regression with Gaussian processes, enabling Markov chain Monte Carlo sampling of posterior distributions.

### 4.1. Comparison with existing measurement methods

The gold standard for determining mechanical properties of skin remains *ex vivo* biaxial testing [25, 17], which provides detailed information about anisotropy and nonlinear stiffening under large deformations. Alternative methods, including uniaxial and bulge testing, are also accurate, but they too require excised tissue [16, 18].

For *in vivo* measurements, current options are limited. The most widely used method is suction-based devices (e.g., Cutometer, CutiScan, Nimble [2, 48, 49]), which apply negative pressure and measure skin deflection. While noninvasive and convenient, these methods primarily yield low-resolution mechanical properties, typically limited to shear modulus estimation. Recent advancements attempt to extract anisotropy information through DIC during suction [50]. Another prominent *in vivo* technique is shear wave elastography, which requires specialized equipment and primarily determines shear modulus, without capturing anisotropic properties [19].

In contrast, our method based on testing with actuated membranes demonstrates the ability to recover not only shear modulus and fiber orientation but also fiber dispersion *κ*, and nonlinear stiffness parameters such as *k*_1_. However, our study reveals that the strains and modes of deformation that can be achieved with actuated membranes attached to the skin does not fully capture nonlinear stiffening effects, particularly the parameter *k*_2_, which is associated with strain-dependent stiffening at larger strains.

Our analysis suggests that the inability to recover *k*_2_ stems from the strain range explored in this study. The influence of *k*_2_ becomes significant only at strains exceeding 10%, whereas the deformations induced by our actuated membrane setup ranges from −5% to 5%. This range is actuation is expected for instance in the case of pre-stressed cosmetic membranes [30]. Suction devices operate at similar strain levels [1], yet they lack the ability to capture anisotropy and nonlinear properties. Shear wave elastography relies on linear analysis of wave speed, providing only shear modulus estimates.

Thus, our study shows that actuated membranes should offer an improvement over current *in vivo* techniques, as they would enable the estimation of anisotropic mechanical properties (*μ, k*_1_, *κ, θ* in the case of HGO-parameterized mechanics), though it still falls short of *ex vivo* biaxial testing.

A key advantage of our approach is the ability to extract mechanical properties without direct force measurements, distinguishing it from suction-based methods. The idea is that membrane designs can be tailored to exert specific forces or strains on the skin, either through magnetic actuation or pre-stressed polymer membranes, both of which have been explored in recent studies [30, 41, 42]. While this approach eliminates the need for force control in favor of membrane design, it still necessitates high-resolution optical measurements such as DIC, which may present practical challenges.

### 4.2. Impact of noise

Our analysis shows that the method is sensitive to noise on the order of 5% or greater in the strain field and with spatially uncorrelated noise (or noise with a small correlation length on the order of 0.1mm). For the small correlation length, even a noise of 1% in strain measurements can add significant uncertainty about nonlinear parameters such as *k*_1_. Beyond 1% strain error, *k*_1_ becomes increasingly more difficult to identify. Nevertheless, the other parameters are still robust to noise, even at 10% noise, as seen in Figure 9. Thus, the proposed method should still work to identify mechanical properties similar to suction and shear wave elastography without force measurements. Fortunately, DIC techniques are well-established and strain errors are typically less than 5% [51, 52]. Therefore, we anticipate that, under best practices, with DIC strain errors < 5%, the method should be able to recover linear and at least some of the nonlinear properties, e.g. *k*_1_ parameter in the case of the HGO model.

### 4.3. Bayesian inference

A potential drawback of our method is the computational cost associated with Bayesian inference. Suction-based and shear wave elastography techniques provide rapid estimates by relying on simple empirical relationships between measured deformation and shear modulus [1]. In contrast, our method involves complex nonlinear deformations, necessitating an inverse finite element analysis. Traditional inverse finite element methods are prohibitively expensive [53], while alternative approaches such as the virtual fields method require problem-specific formulation [54]. In combination, sampling posteriors via MCMC limits the speed with which predictions can be made. Running multiple chains can take hours to converge for a single data point, preventing real time feedback. Other methods to sample posteriors can be significantly faster, such as variational inference [55].

Given the inherent ill-posedness of the problem, Bayesian inference is preferable over deterministic inverse methods, as it accounts for uncertainty in parameter estimation [56]. However, the high computational burden of full forward simulations necessitates model reduction techniques. We employed PCA for dimensionality reduction and GP regression for surrogate modeling [9]. Alternatives include artificial neural networks for the nonlinear regression task [57]. Future work will also explore alternative dimensionality reduction techniques, for example nonlinear manifold reconstruction [58]. One area of focus will be robustness against noise [59].

### 4.4. Limitations

The present study is based on synthetic data. The rationale is to test numerically whether deformation fields induced by active membranes can be used as a method for material identification. Thus, the methodology and numerical studies shown here are a necessary pre-requisite for future experimental work. We showed that indeed, existing prestressed membranes [30] or actuated membranes (for example magnetic actuation [42]) can be used to determine skin material properties. This work provides confidence in ongoing experimental efforts to demonstrate the practical use of this methodology in the near future.

Another limitation of our study is its reliance on a specific material model—the Holzapfel-Gasser-Ogden (HGO) hyperelastic model [39]— which may not be the exact material model of skin across the human population. While HGO is well-suited for mouse, porcine, and human skin, based on existing data [60, 17, 25], future investigations should focus on more general, perhaps data-driven alternatives [61].

## 5. Conclusion

Overall, our study establishes a theoretical and numerical foundation for improving *in vivo* skin mechanics measurement. While further experimental validation is required, our findings suggest that actuated membranes present a promising alternative to quantify nonlinear hyperelastic skin properties compared to traditional methods such as suction or shear wave elastography. Key characteristics of the method are the Bayesian approach, which can effectively deal with the ill-posed problem, especially in the presence of noise. Another key idea is to invert the strain field to obtain mechanical properties without force data, relying instead on known properties of an actuated membrane attached to the skin. Lastly, the fact that the method is Bayesian is enabled by replacing the finite element solver with a Gaussian process surrogate trained on finite element simulation data projected onto a reduced basis.

## Data Availability

A GitHub repository with the code used to run all experiments is available at https://github.com/mwilkinson1/bayesian-active-membranes

## Acknowledgements

This work was supported by the National Institute of Arthritis and Musculoskeletal and Skin Diseases of the National Institute of Health under award R01AR074525 to Adrian Buganza and by the National Institute of General Medical Sciences under award R21GM137275 to Adrian Buganza and Andres Arrieta.

